# Using Latent Chemical Recognition in an Evolved Periplasmic Binding Protein Family to Diversify Biosensors

**DOI:** 10.64898/2026.08.04.742673

**Authors:** Eryney Marrogi, Aaron Nichols, Henry Lester, Anand Muthusamy

## Abstract

Genetically encoded fluorescent biosensors have gained traction in neuroscience as continuous, reagentless reporters of cellular activity *in situ*. These sensors, often soluble, can also provide time-resolution in multiple biosensing form factors: benchtop and wearable devices, lyophilized powder tests, and smartphone-based diagnostic tests. These biosensors often take advantage of naturally occurring conformation-switching but require extensive screening to optimize the linkers to a fluorescent reporter. Each new target ligand often requires its own engineering campaign. We asked whether sensors evolved towards a particular target retain useful recognition scope for others. We screened a family of 18 OpuBC-cpGFP sensors evolved toward nicotinic agonists, SSRIs, opioids, and other neural drugs, against 63 structurally diverse compounds. We found that 24 ligands activated at least one biosensor with ΔF/F_0_ > 0.3, sufficient to begin directed evolution, with 8 of those ligands activating at least one biosensor with ΔF/F_0_ > 1.0, the regime of dynamic range usable in end applications. With 124 ligand-biosensor pairs in total, we found multiple leads suitable for directed evolution. Most notably, ligands participating in hits spanned well beyond neural drugs and included DEHP, ergothioneine, ciprofloxacin, thiamine, betahistine, L-carnitine, and L-thyroxine. Across the biosensor family, mutation distance weakly predicted substrate scope. In particular, we observed sequence-function cliffs that could be exploited for future protein engineering campaigns. Thus, broad screening of performant scaffolds offers rapid bootstrapping in biosensor engineering particularly for exogenous molecules.

## Introduction

Protein engineering is increasingly informed by computational models that predict protein structure, generate sequences constrained to specified backbones, and design new folds or binding interfaces towards novel functions^1–4^. However, structural prediction alone underdetermines protein function, and in these cases large experimental datasets are required to shore up computational methods. Genome editing via engineered proteases, or viral capsid tropism can be steered with large scale mutational scans. We lack high throughput biophysical assays for engineering conformational changes related to molecular recognition and allosteric coupling in new scaffolds. Complicating the task, ligand binding and conformational change must be evaluated for specific trajectories of movement , responsiveness in physiological environments, and selectivity. Furthermore, for a biosensor, this must also produce a measurable output that can inform additional protein engineering campaigns. The broader application of such approaches depends on experimental systems that translate the desired molecular function as predicted in the computational design stage into a quantified phenotype.

Biological signaling frequently depends on ligand-induced changes in protein conformations, dynamics or intermolecular interactions. These mechanisms underlie the operation of receptors, transport proteins and allosteric enzymes, and are also exploited in the construction of genetically encoded biosensors. In these systems, a ligand-recognition event is coupled to an electrochemical or fluorescent reporter that allows molecular concentrations to be measured continuously without the addition of exogenous reagents, and to do so in ideal physiologic conditions.^5–9^ Recent demonstrations show that machine-learning-designed binding domains can be incorporated into artificial allosteric switches, which suggests that computationally designed recognition elements can be coupled to reporter proteins^9^. Nevertheless, successful construction of these systems requires experimental identification of architectures in which binding and reporter activity are accurately coupled to optimally capture the value of scaled AI-mediated design. Recent work showing that proteases redesigned with AI provide improved starting points for subsequent laboratory evolution illustrates the value of combining machine guided design with a tightly coupled functional selection^10^.

We asked if periplasmic binding protein (PBP) biosensors could provide a scaffold for a similar protein engineering task. PBPs generally contain two lobes that close around a ligand in a “Venus flytrap” motion (**Fig 1a, b**). Insertion of a circularly permuted green fluorescent protein (cpGFP) into an allosterically responsive region of the PBP via linkers can convert this ligand-dependent motion into a change in fluorescence.^5^ The resulting genetically encoded sensor can be purified for analytical assays, incorporated into experimental devices, or expressed in cells and subcellular compartments for monitoring of physiologic processes^6–8^. They therefore provide both a ligand-binding scaffold and an intrinsic quantitative readout, properties that are useful for engineering sensors intended for continuous molecular monitoring. The development of PBP-cpGFP sensors requires identification of a suitable binding protein, selection of reporter insertion sites and linkers, modification of the binding pocket, and parallel optimization of affinity, selectivity, expression and fluorescence dynamic range. Directed evolution can introduce new ligand contacts while retaining alternative recognition modes, and intermediate constructs may possess functional profiles that are absent from the final optimized sensor. Systematic screening of an evolved biosensor lineage can identify latent recognition of non-target substrates and generate experimentally measured sequence-function data for subsequent computational or directed-evolution campaigns. This strategy begins from scaffolds that already produce appreciable output and focuses design on improving an observed response rather than simultaneously creating ligand recognition and signal transduction *de novo*.

**Figure 1.**
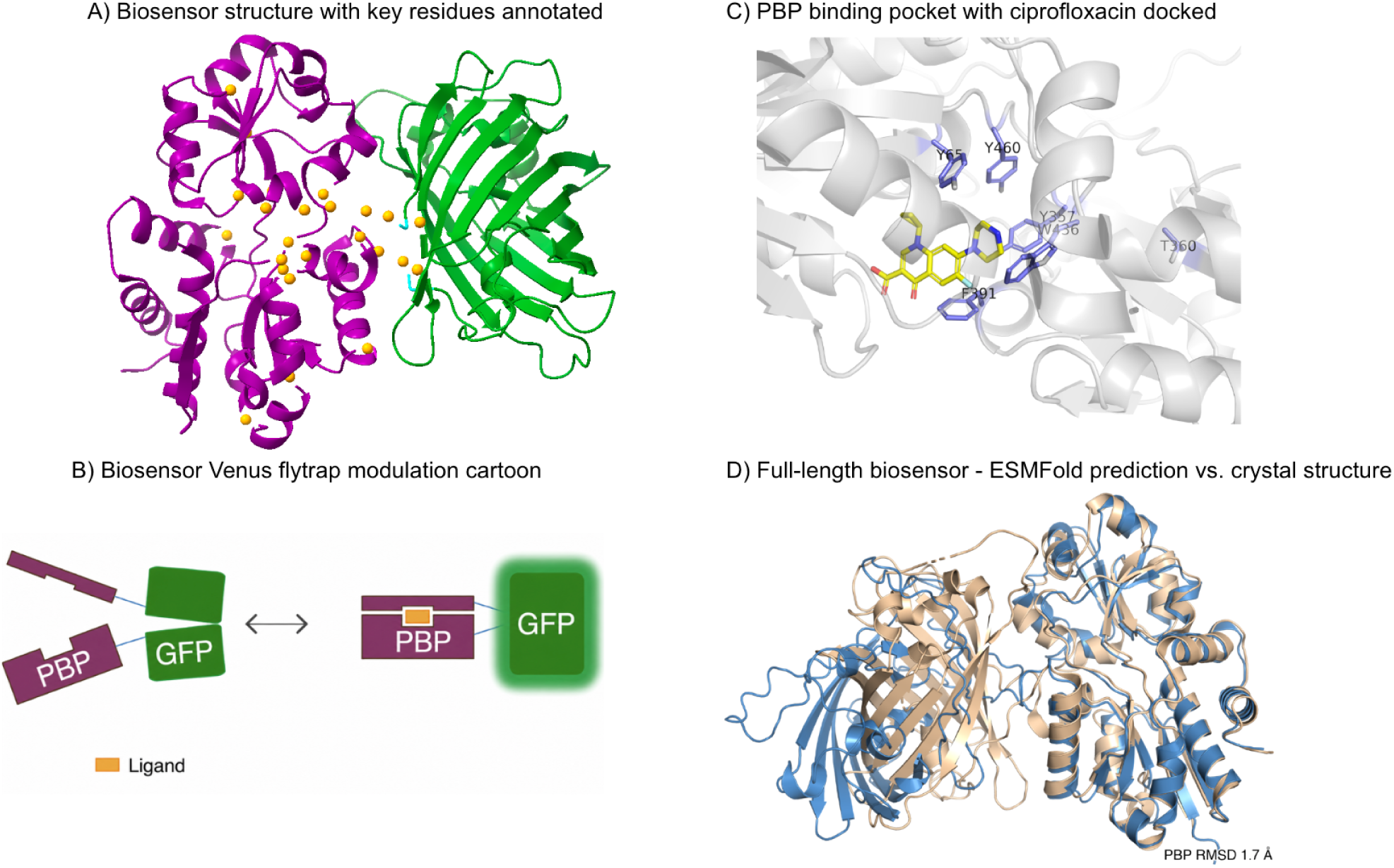
The OpuBC-cpGFP biosensor scaffold and its ligand-binding pocket. (A) Crystal structure of iNicSnFR3a (cc93; PDB 7S7U^8^) composed of the mutant OpuBC (magenta), GFP (green), and the connecting linkers (cyan). Orange spheres mark the 31 Cα positions that vary across the 17-member scaffold family, the four aromatic-cage residues Y65, Y357, F391, and W436 are drawn as yellow sticks, and the GFP chromophore is shown in limon. (B) Sensing mechanism where the PBP closing on its ligand pulls a linker glutamate out of the cpGFP, relieving quenching of its fluorophore, thereby increasing its brightness. (C) Binding pocket with ciprofloxacin (yellow sticks) placed by blind diffusion docking (DiffDock-L).^15^ Aromatic-cage and pocket residues (Y65, Y357, F391, W436, Y460, T360) are shown as marine sticks (D) Full-length structure prediction (ESMFold^2^, mean pLDDT ≈ 70) superposed on the 7S7U crystal (wheat) over the PBP domain. The predicted PBP lobes (blue) overlay the crystal closely (Cα RMSD 1.7 Å over the folded core), whereas the predicted linker and cpGFP module are displaced by ≈ 21 Å. Generally, we find that AI-based structure prediction poorly recapitulates the linker and cpGFP position relative to the PBP, missing the transduction mechanism.

This study examines latent chemical recognition within a family of OpuBC-cpGFP biosensors. OpuBC is a choline-binding PBP^11^ that has been repurposed through protein engineering to detect nicotine, nicotinic agonists, selective serotonin reuptake inhibitors (SSRIs), opioids, and other drug-like molecules where pharmacokinetic monitoring is useful^8,12–14^. We screened 18 biosensors from this family against 63 structurally diverse compounds selected for an interest in human and environmental health monitoring, spanning metabolites, pharmaceuticals, hormones, neurotransmitters, and environmental chemicals. We define “latent recognition scope” to denote detection of ligands outside of a scaffold’s anticipated natural and engineered targets, “ligand hit breadth” to denote the number of sensors activated by a ligand, and “sensor hit breadth” to denote the number of ligands detected by a sensor. We asked whether sensors evolved for specific neural drugs retained measurable responses to ligands outside their intended recognition classes, whether small numbers of mutations produced predictable changes in ligand scope, and whether any biosensors could serve as leads for potential applications.

## Results

### Broad screening reveals latent recognition

The panel comprised 17 OpuBC-cpGFP scaffold sensors plus one null biosensor, iFentanylSnFR2 W436L (a fentanyl biosensor with a key aromatic residue mutated, producing zero response to its ligand). The primary lineage runs from early nicotine-responsive constructs through iNicSnFR3a (cc93), iNicSnFR3b (v7), later acetylcholine intermediates, and iAChSnFR (v9), with serotonin, escitalopram, cytisine, fluoxetine, tapentadol, and levorphanol facing branches. Across the 17 scaffolds, pairwise identity ranges from 95.0 to 99.8%. Only 31 of 521 aligned positions vary, and several positions lie near residues implicated in selectivity, including Y357 and W436. W436 lies in the aromatic cage that recognizes cationic ligands in this family^16^, which makes mutations at this position plausible switches for substrate scope (**Figure 1C, D**). The sensors are highly similar in sequence but functionally distinct in terms of substrate recognition, suggesting that even point mutation screens can probe the potential for re-targeting the PBP.

Screening 18 sensors against 63 ligands at a single concentration, we computed ΔF/F_0_ relative to buffer controls, with responses ranging from −0.90 to 12.1. We defined biosensor-ligand pair responses of ΔF/F_0_ > 0.3 as “engineerable” and ΔF/F_0_ > 1.0 as a “strong response” suggesting the potential for immediate use in quantitative applications. 24 of 63 ligands activated at least one sensor above 0.3, and eight exceeded 1.0 including nicotine, thiamine, DEHP, ergothioneine, betahistine, ciprofloxacin, L-carnitine, and L-thyroxine.

The pair-level screen was sparse but we observed at least one engineerable hit across most ligands. Of 1,134 sensor-ligand combinations, 124 exceeded 0.3 and 50 exceeded 1.0. Nicotine gave the largest native-class signal (maximum ΔF/F_0_ = 12.1, with 10 sensors above 1.0). This result was expected as many of the tested sensors derive from an engineered family of nicotine sensors that we previously reported. Several non-native ligands also produced large responses in some biosensors with ΔF/F_0_ of 4.72 for thiamine, 3.24 for DEHP, 3.04 for ergothioneine, 2.75 for betahistine, and 2.50 for ciprofloxacin.

We found hits in all three compound classes screened: “alkaloid-like”, “non-alkaloid charged”, and “non-alkaloid, uncharged”. The charged non-alkaloid group was the smallest in count but yielded three strong responses of six total compounds. The larger uncharged non-alkaloid group presented hits for DEHP, ciprofloxacin, and L-thyroxine. **Figure 2C** shows representative structures spanning the charge, size, and shape range. The central observation is that a family founded on a choline-binding PBP can be exploited for promiscuous binding to compounds of widely varied polarity and shape.

**Figure 2.**
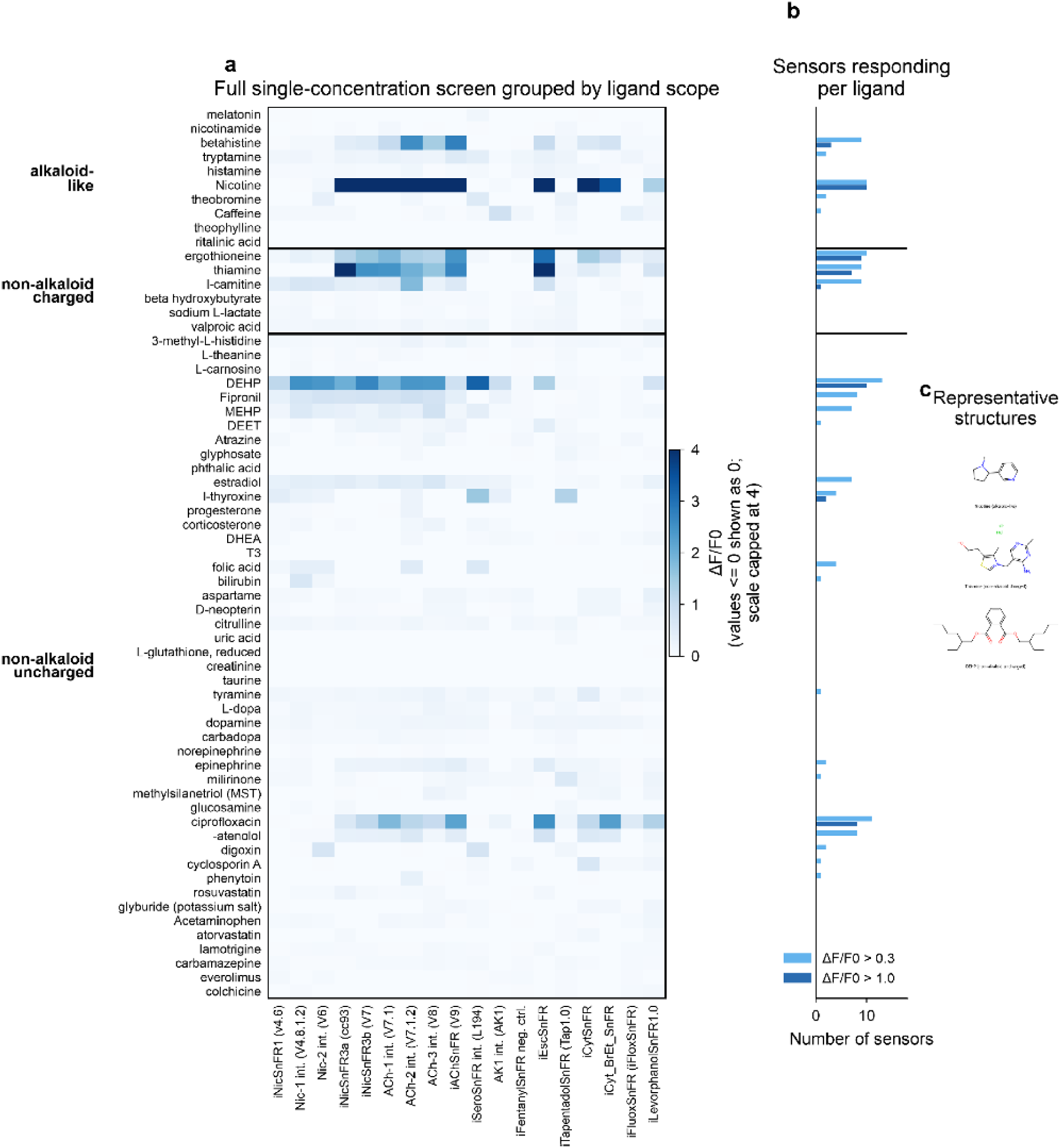
Compound screen grouped by ligand class. (A) Full 63 ligand × 18 sensor response matrix, grouped into alkaloid-like, non-alkaloid charged, and non-alkaloid uncharged ligands. Values ≤ 0 are displayed as zero and the color scale is capped at ΔF/F_0_ = 4.0 to visualize weak and moderate hits. The largest overall response belongs to nicotine binding to one of its biosensors with response ΔF/F_0_ = 12.1. (B) Number of sensors responding to each ligand at ΔF/F_0_ > 0.3 and > 1.0. (C) Representative structures from the 3 scope classes, showing nicotine, thiamine, and DEHP.

Ergothioneine and thiamine exceeded 1.0 in 9 and 7 sensors, respectively. DEHP presented the broadest non-nicotinic hit, activating 13 sensors above 0.3 and 10 above 1.0, and ciprofloxacin activated 11 and 8, respectively. MEHP, which differs from DEHP only by replacement of one ethylhexyl ester with a carboxylate, activated no sensor above 1.0, suggesting general recognition of a particular hydrophobic shape rather than a generic phthalate response. Notably, several sensors did not respond to DEHP, suggesting there is not a generic, non-specific interaction between it and the linkers and cpGFP.

### Sequence-function cliffs reshape ligand fingerprints

Mutation distance predicted functional similarity only weakly. Across the 17 scaffolds, pairwise mutation count correlated negatively with response-profile correlation (r = −0.364, p = 1.4 × 10^-5^), yet the spread around this trend was large, and many close neighbors displayed highly varied fingerprints. Through parent-child steps of 1 to 3 mutations, 12 ligands crossed the meaningful-response threshold (**Figure 3B**). Tap1.0 and v7 illustrate the extreme case of this phenomenon. These two sensors differ by a single internal mutation, W436A, yet 12 ligand-response states changed; Tap1.0 gained 2 hits and lost 10 relative to v7, and their response fingerprints were uncorrelated (r = −0.017). Two further examples reinforce the pattern: Tap1.0 and v8 differ by 3 internal mutations (r = −0.026), and iEscSnFR and Tap1.0 differ by two (r = 0.002). We call these sequence-function cliffs, in which a small sequence change produces a large functional change, a key exploit for future screening and machine-guided protein engineering campaigns.

**Figure 3.**
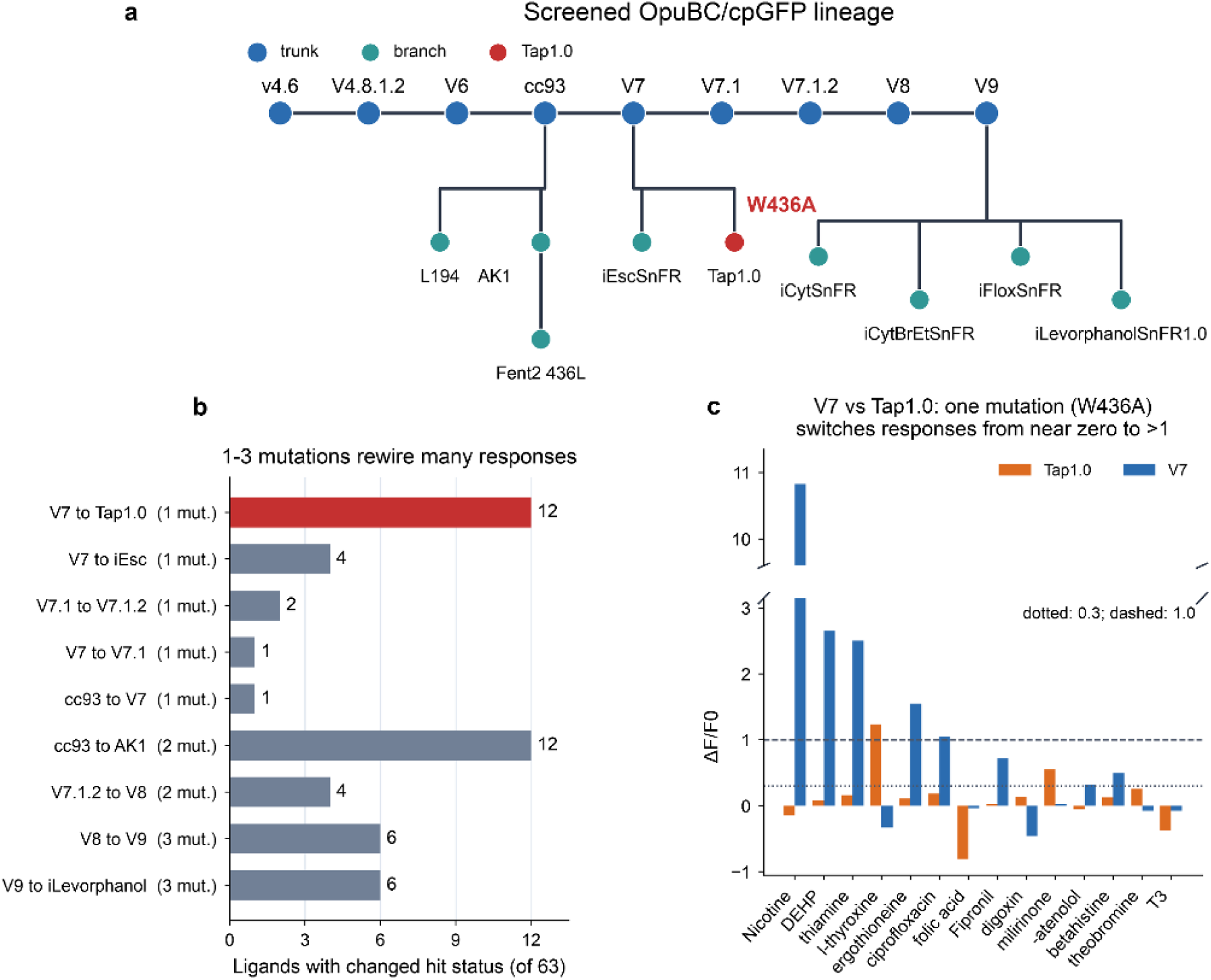
A few mutations can redirect ligand recognition across the OpuBC-cpGFP family. (A) Screened sensor lineage in the simplified paper-facing taxonomy. Tap1.0 is highlighted because it differs from v7 by W436A, and iFentanylSnFR2 436L is shown as the sequence-outgroup negative control but excluded from scaffold-only sequence analyses. (B) Number of ligands changing hit status across each parent-child step of 1 to 3 mutations, where a status change means crossing ΔF/F_0_ = 0.3 in either direction. v7→Tap1.0 and cc93→AK1 each changed 12 of 63 ligand-response states, after 1 and 2 mutations respectively. (C) Tap1.0 and v7 responses for the 14 ligands with the largest absolute differences. The broken y-axis isolates the nicotine response (ΔF/F_0_ = 10.8 in v7) while preserving the scale of the remaining ligands. Dotted and dashed lines denote ΔF/F_0_ = 0.3 and 1.0.

### Ciprofloxacin sensors approach an application-relevant range

We fit Hill curves for selected hits to further characterize ligand-sensor pairs. Four sensors gave clear dose-dependent ciprofloxacin responses with acceptable fit quality. iEscSnFR, v7.1, v9, and v7 gave EC_50_ values of 22.4, 35.9, 45.3, and 53.7 µM (r^2^ = 0.990 to 0.999). These EC_50_ values exceed standard plasma exposure, yet the curves already produce measurable signals within the clinically relevant range. The ciprofloxacin label reports a steady-state C_max_ of 2.97 µg/mL for 500 mg administered orally every 12 hours, which corresponds to approximately 9 µM.^17^ The four sensors displayed a response of ΔF/F_0_ of 0.45 to 1.67 at 6.3 to 20 µM ciprofloxacin. We anticipate that a focused campaign evolving the biosensor for ciprofloxacin should improve both affinity and slope to quickly yield a sensor with classical Hill fit.

Other ligands showed substantive responses encouraging directed evolution toward their application. For example, thiamine, ergothioneine, and L-carnitine produced appreciable responses into the low-µM range but they did not reach saturation in our dose responses. Thus their likely EC_50_s are above physiological plasma ranges.

## Discussion

Our study demonstrates that an evolved PBP-cpGFP biosensor family has considerable latent recognition capacity beyond its original, planned targets. Screening 1,134 sensor-ligand combinations identified 124 responses above ΔF/F_0_ = 0.3 and 50 above ΔF/F_0_ = 1.0. Most notably, these ligands were structurally and chemically distinct from the cationic neural drugs that the original family was designed to recognize, including ciprofloxacin, DEHP, ergothioneine, thiamine, betahistine, L-carnitine, and L-thyroxine. In essence, our argument is four-fold: (1) good scaffolds are rare, (2) we have previously found PBPs to be highly tolerant of mutations, (3) an expeditious directed evolution campaign will seek to push promiscuity to its limit, and (4) directed evolution and AI-guided design can take over from that point to “hill-climb” to performant mutants.

Screening methods as in this work produce useful starting points for further directed evolution campaigns. These endeavors can begin by optimizing an observed, mechanistically coupled response rather than independently identifying a binder, establishing a ligand-dependent structural transition, and discovering a reporter architecture capable of detecting that transition. This is critical for analytes without known natural receptors or for those with binding proteins that have been categorized nominally as an “X-binding protein.” Further studies are required to ensure apparent hits are not the result of pH artifacts, aggregation, or indirect modulation of the biosensor. While these spurious signals are rare in our screens, particularly with buffering in 3x PBS and visual inspection of plates prior to each experiment, they are possible and would be revealed in follow-up mechanistic studies and directed evolution.

The long-term objective of computational biosensor design is to generate recognition elements whose binding properties and output mechanisms are specified jointly, such that the design stage can be more grounded in physiologic reality. Such proteins could directly modulate reporters such as fluorescent proteins, enzymes, and nanopores. Current design methods can generate candidate motifs and binding interfaces, and ML-designed receptors have recently been incorporated into synthetic allosteric switches^9^. However, joint design remains a challenge as specificity, affinity, expression and stability must be satisfied simultaneously to contribute meaningfully to design. The displacement of the predicted cpGFP module relative to the experimental structure in **Figure 1D** illustrates this limitation of computational prediction of allosteric coupling.

Our findings suggest a pragmatic route for tying computational protein design to quantitative functional selection. Reporter-coupled scaffolds can first be screened to identify proteins in which a target ligand already produces a measurable output. Computational models can then search locally around these experimentally validated starting points for mutations that improve ligand recognition domains towards selectivity while preserving the existing allosteric architecture. Within the context of biosensors, the fluorescent output supplied the functional phenotype required to compare proposed designs directly in an iterative manner.

The sequence-function cliffs described in our study across the OpuBC-cpGFP lineage further supports this approach. Closely related sensors frequently exhibited divergent ligand-response profiles, and individual substitutions could substantially redirect chemical recognition. As an example, the single substitution separating v7 and Tap1.0 led to variations in 12 ligand-response states and reduced the correlation between their binding profiles to virtually zero. Similarly, the single T360S substitution that separates v7 and iEscSnFR was associated with a 2.4-fold improvement in apparent ciprofloxacin sensitivity. These observations indicate that narrow mutational changes can generate substantial functional diversity that can then be captured quickly in broad ligand screening experiments that expose this diversity, as was seen in our study.

Taken together, these results establish that evolved allosteric biosensor families can serve as starting libraries for computational protein engineering. Their value derives from the presence of molecular recognition, conformational coupling and quantitative readout within a single protein. This strategy reduces the time required to develop sensors for molecules lacking established recognition scaffolds by identifying latent responses and exploiting their large functional changes through small numbers of mutations. This ultimately provides a means of linking computationally proposed protein sequences to experimentally measured functions, addressing a central limitation in the application of machine-guided design of allosteric interactions. Creating new biosensors, particularly those based on PBPs, enables the same molecule to serve in cell culture, *in vivo* animal, point of care, wearable device, and lyophilized powder portable field tests^18,19^. Furthermore, we anticipate this coupled screening/design approach will be necessary to generate sufficient quantities of data to train models for allostery prediction in the future.

## Methods

### Sensor panel and ligand screen

The dataset comprises 18 genetically encoded fluorescent biosensors screened against 63 ligands plus a buffer control. Sensor names, protein sequences, ligand names, SMILES strings, and single-concentration responses were parsed and analyzed using our published screening workflow for generating CNS drug biosensors.^20^

Compounds were purchased from Sigma Aldrich, Thermo Fisher, and Tocris, stored under the manufacturer’s recommended conditions, and used to prepare solutions for screening. Each compound was prepared in a 2 mM stock solution, maximizing the aqueous fraction where possible. We solubilized compounds in 3x PBS pH 7.4 and cut in organic solvent as required; some compounds were insoluble in aqueous buffer and 100% organic solvent was used (typically DMSO or ethanol). We prepared matching buffers as negative controls (no ligand) to match each compound’s associated buffer as some biosensors show minor responses to organic solvents. The 2 mM stocks were used in serial dilutions to prepare solution plates for screening.

Compound solutions were mixed into purified biosensor solutions (111 nM) to give 100 nM final biosensor concentration with compound concentrations ranging from 200 µM to 63.3 nM. We used an epMotion 5075 liquid handling robot for this step and included aspirations for mixing three times after adding the compound. The plates were then read in a Tecan Spark 10M for fluorescences measurements (485 nm excitation, 20 nm bandwidth; 535 emission, 25 nm bandwidth set for GFP).

We calculated ΔF/F_0_ for responses for each biosensor, setting F_0_ as that biosensor’s buffer-matched baseline in the absence of any ligand. Screening SEM values were propagated from the ligand and matched baseline SEM values and are reported in the supplement. We defined a “hit” as a biosensor-ligand pair’s response of ΔF/F_0_ > 0.3 and a “strong hit” as ΔF/F_0_ > 1.0. Ligand hit breadth and sensor hit breadth were computed as the number of sensors or ligands exceeding that threshold.

### Ligand-scope classification

Ligands were assigned to 3 classes for the analysis in **Figure 2**: “alkaloid-like”, “non-alkaloid, charged”, and “non-alkaloid, uncharged”. Alkaloid-like ligands included nicotine and related nitrogenous bioactive compounds. Charged compounds, including zwitterions and permanent cations, were identified from formal charges assigned in RDKit-parsed SMILES^21^, and the remainder of the ligands were assigned to the non-alkaloid uncharged class.

### Dose-response fitting

Dose-response files were extracted from the raw plate-reader Excel workbooks (Tecan Spark 10M). Each experiment contained 3 replicate rows, a buffer baseline column, and 8 ligand concentrations beginning at 200 µM and serially diluted by a factor of √10. ΔF/F_0_ was computed from the mean ligand and mean baseline fluorescence, and standard error of the mean (SEM) was propagated across the baseline and replicate wells. Four weak or negative extra curves were excluded from the processed set: MEHP_cc93, aspartame_iLevaSnFR, bilirubin_V4.8.1.2, and theobromine_Tap1.0. In the ciprofloxacin plates, well H2 at 200 µM was excluded because the liquid-handling robot failed to dispense into that well. For that dose response, the 200 µM point was averaged over the remaining two replicates.

Curves were fit with a 4-parameter Hill model, F(x) = baseline + ΔFmax / (1 + (EC50 / x)^n^). Fits used the SciPy curve_fit routine^22^ with multiple starting EC_50_ and Hill coefficients, with lower bounds of ΔF_max_ ≥ 0, EC_50_ ≥ 1 × 10^-4^ µM, and n ≥ 0.1, and upper bounds of EC_50_ ≤ 1 × 10^6^ µM and n ≤ 10. Curves with a response range below ΔF/F_0_ of ∼0.1 were flagged as noise. Fits with EC_50_ < 400 µM and ΔF_max_ < 20 were treated as reasonable and acceptable, and the remainder were treated as out-of-range. For **Figure 4**, ciprofloxacin curves were highlighted as a potential application-ready biosensor because 4 sensors gave monotonic dose-dependent responses with measurable signals from the single-digit to low-tens of micromolar range and R^2^ > 0.99.

**Figure 4.**
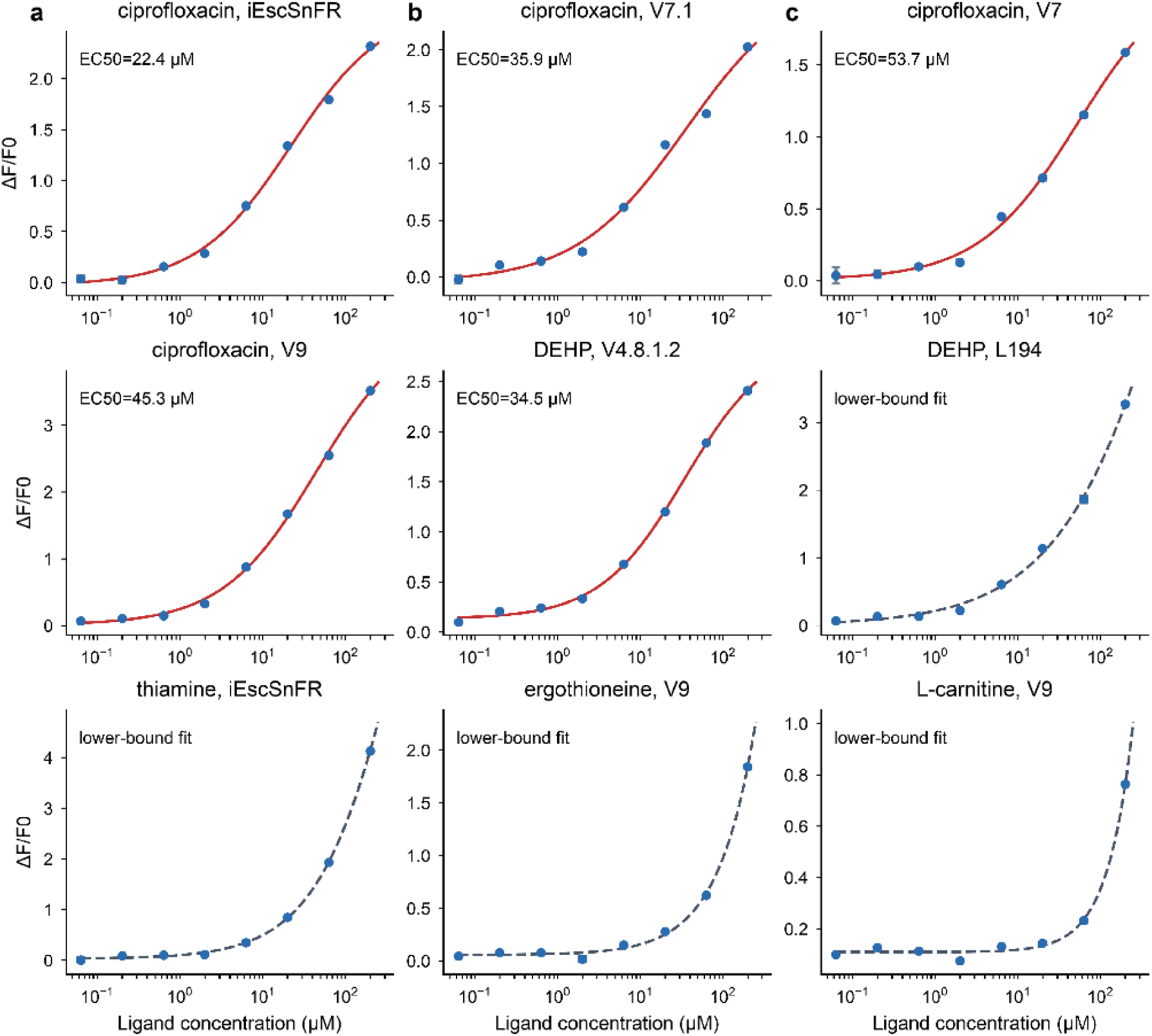
Dose responses for the strongest latent-recognition leads from our screen. Points show mean ΔF/F_0_ from three replicates, and error bars show propagated SEM. Ciprofloxacin responses in iEscSnFR, v7.1, v7, and v9 give displayed EC_50_ values of 22.4 to 53.7 µM after excluding the known bad H2 well at 200 µM and recomputing that point from the remaining 2 replicates. Solid red curves indicate good fits, and dashed gray curves indicate fits for dose responses that did not reach near-saturation. DEHP includes a good fit for v4.8.1.2 and non-saturation with L194. Thiamine, ergothioneine, and L-carnitine show large high concentration responses but need affinity maturation for most physiological applications.

### Sequence-function analysis

Protein sequences were taken from the curated sensor sequence table reported in Supplementary Table S7. Pairwise mutation events were counted across the 17 OpuBC/cpGFP scaffold sensors after exclusion of the negative-control outgroup iFentanylSnFR2 436L. A terminal one-residue length difference was ignored for the Tap1.0/v7 comparison, whose internal sequence comparison identifies a single W436A difference. Functional similarity was computed as the Pearson correlation between full ligand-response profiles. For **Figure 3B**, parent-child steps were taken from the simplified lineage in **Figure 3A** and restricted to pairs separated by 1 to 3 mutations. Each ligand was classified as a hit or non-hit across the ΔF/F_0_ > 0.3 threshold in parent and child, and ligands crossing that threshold in either direction were counted as changed states.

## Data and Code Availability

All processed data, analysis code, and figures are publicly available at https://github.com/eryney/biosensor-latent-recognition. The repository contains the processed single-concentration response and propagated SEM matrices, the 49 processed dose-response curves and their Hill fits, ligand SMILES and scope classifications, sensor sequences, the supplementary tables workbook, and the final figures. The analysis pipeline is included: analysis/make_figures.py regenerates Figures 2–5 from the processed tables, analysis/compose_figure1.py assembles Figure 1 from the rendered structural panels, and analysis/build_supplement_tables.py rebuilds the supplementary workbook. The raw plate-reader workbooks are available from the authors on request. Code is released under the MIT License and data under CC BY 4.0.

## Notes

### Competing Interest Statement

The authors have declared no competing interest.

